# A Fully-Adjusted Two-Stage Procedure for Rank Normalization in Genetic Association Studies

**DOI:** 10.1101/344770

**Authors:** Tamar Sofer, Xiuwen Zheng, Stephanie M. Gogarten, Cecelia A. Laurie, Kelsey Grinde, John R. Shaffer, Dmitry Shungin, Jeffrey R. O’Connell, Ramon A. Durazo-Arvizo, Laura Raffield, Leslie Lange, Solomon Musani, Ramachandran S. Vasan, L. Adrienne Cupples, Alexander P. Reiner, Cathy C. Laurie, Kenneth M. Rice

## Abstract

When testing genotype-phenotype associations using linear regression, departure of the trait distribution from normality can impact both Type I error rate control and statistical power, with worse consequences for rarer variants. While it has been shown that applying a rank-normalization transformation to trait values before testing may improve these statistical properties, the factor driving them is not the trait distribution itself, but its residual distribution after regression on both covariates and genotype. Because genotype is expected to have a small effect (if any) investigators now routinely use a two-stage method, in which they first regress the trait on covariates, obtain residuals, rank-normalize them, and then secondly use the rank-normalized residuals in association analysis with the genotypes. Potential confounding signals are assumed to be removed at the first stage, so in practice no further adjustment is done in the second stage. Here, we show that this widely-used approach can lead to tests with undesirable statistical properties, due to both a combination of a mis-specified mean-variance relationship, and remaining covariate associations between the rank-normalized residuals and genotypes. We demonstrate these properties theoretically, and also in applications to genome-wide and whole-genome sequencing association studies. We further propose and evaluate an alternative fully-adjusted two-stage approach that adjusts for covariates both when residuals are obtained, and in the subsequent association test. This method can reduce excess Type I errors and improve statistical power.

## Introduction

Linear regression-based tests of associations of genetic variants with a quantitative trait can be sensitive to departure of the trait distribution from normality, particularly when testing rare variants. To address this problem, an approach that is widely used in genetic association studies (within the regression framework) is to apply a rank-normalization of trait values, followed by subsequent analysis of the rank-normalized trait as the analysis outcome (see e.g. (ref. 1, 2) and a comprehensive review in (ref.3)). In the conte*x*t of rare variants, it was shown by Tang and Lin (ref.4) that applying rank-normalization on traits prior to any analysis and testing helps to control the rate of Type I errors and increase statistical power. However, the factor actually determining the statistical properties of regression-based trait-variant association tests is not the distribution of the trait but instead its distribution after regressing out covariates. Indeed, it was shown in (ref.3) that rank-normalizing traits may still result in non-normal residuals, resulting in invalid Type I error rate control, with the problem being most severe when the distribution of the residuals is heavily skewed. Recent Genome-Wide Association Studies (GWASs) have instead used a different approach, applying the rank-normalization to the residuals that were generated by regressing the trait on covariates (ref.5, 6–9) in stage 1, and then using these transformed residuals as the outcomes in subsequent analyses, without further adjustment for covariates (stage 2). For GWASs, which primarily address the analysis of common genetic variants, this partly-adjusted two-stage approach has been criticized (ref.10, 11) due to potential loss of power, and biased estimates when covariates are correlated with genotypes. However, some researchers (ref.12) still suggest that this approach is appropriate for analysis of rare variants. As rare variant analysis is the focus of most Whole Genome Sequencing (WGS) studies, which are currently underway, there is now a strong motivation to better understand why these problems occur and how transformations and covariate associations interplay to affect them, and to provide a comprehensive framework for genetic association analyses for quantitative traits that is appropriate under a wide range of settings.

In this investigation, we propose a fully-adjusted two-stage approach that both provides the protection of rank-normal transformations, and also mitigates the potential for mis-calibrated inference. In Stage 1 we regress the trait on covariates and obtain residuals, which we rank-normalize. In Stage 2 we use these rank-normalized residuals in association analysis with the genotypes, but adjusting for the same covariates used in Stage 1. The covariate adjustment at Stage 2 differentiates the proposed method from the commonly-used partly-adjusted two-stage approach. Here, we show that using the partly-adjusted method can lead to (1) invalid estimates of regression coefficients’ standard errors, and invalid Wald and Score tests; and (2) residual confounding, because rank-normalization interferes with the adjustment for covariate effects. Adjustment for covariates in stage 2 alleviates both of these issues. Surprisingly our approach may increase statistical power compared to the partly-adjusted approach even if the residuals are perfectly Normally distributed. We investigate our approach from two perspectives. First, we separate the issues of covariate adjustment and rank normalization and study, via linear regression theory, the effects of covariate adjustment alone (or lack thereof) on testing associations with the residuals in the absence of rank normalization. Second, we perform simulations mimicking the settings used in (ref.12), comparing approaches for rank-normalization and covariate adjustment of traits and residuals under normality and deviations from normality, and by the strength of confounding effects. Finally, we demonstrate the undesirable consequences of using rank-normalized residuals in genetic association testing without covariate adjustment in two applications: multiple GWASs in the Hispanic Community Health Study/Study of Latinos, and a WGS analysis of blood hemoglobin levels using three studies from the Trans-Omic Precision Medicine (TOPMed) TOPMed phase 1: the Framingham Heart Study (FHS), Jackson Heart Study (JHS) and Old Order Amish Study (Amish). We show how our fully adjusted two-stage approach addresses these problems.

## Materials and Methods

### Linear regression and residuals

For person *i* we assume that a linear regression model holds, in which each individual has quantitative trait *y_i_*, a *p* × 1 covariate vector ***x_i_*** and a *k* × 1 vector of genotypes ***g_i_*** *(in single variant testing k* is equal to 1). For simplicity, our derivations assume that the observations are independent with identically distributed (iid) error terms; however, they extend straightforwardly to correlated outcomes, for instance when there are genetically related individuals modelled via a kinship or genetic relatedness matrix.

According to the linear regression model:
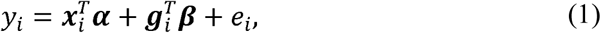

for covariate effects ***α*** and genetic effects ***β***, and where the error terms *e_i_*, *i* = 1,…*,n* are independent and identically distributed (iid). In GWAS and WGS analyses the main focus is testing the null hypothesis *H*_0_:***β*** = **0**. This can be done by first obtaining ordinary least squares estimate 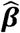 (in addition to 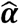) and a corresponding standard error estimate 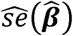, and then using these in a Wald test. A popular alternative is the Score test, which does not rely on estimating ***β*** (ref.13), but rather, is based on a model that only estimates ***α*** via a ‘null model’, in which the outcome *y_i_* is regressed on just the covariates *x_i_*. Residuals are obtained from this null model, and their association with genotype is then assessed. We denote these residuals by **ϵ** = (ϵ_1_,…,*ϵ_n_*)*^T^*, where 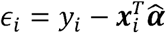. A commonly used Score test in genetic association studies that interrogate rare variants, is the Sequence Kernel Association Test (SKAT, (ref.14)), which tests the association of a set of rare variants with the outcome.

### Two-stage approaches for genetic association analysis

In the first stage of a two-stage approach, the null model is fit, the residual vector **ϵ** calculated, and a rank-normalizing transformation is applied to **ϵ**. This means that entries of **ϵ** are matched with quantiles of the normal distribution, so that the transformed values maintain the same order (or rank) as the original residuals, but follow the normal *N*(0,1) distribution (ref.4). In the second stage the transformed residuals, which we denote *RN*(**ϵ**), are tested for association with the genotypes. It is common in practice to leave this second stage unadjusted for covariates, but in our fully adjusted approach we do adjust for them. This means that the design matrix used when testing a genotype in the second stage differs between the partly- and fully-adjusted approaches. In the partly adjusted approach, the design matrix consists of only an intercept (in addition to the tested genotype), while in the fully adjusted approach, the design matrix is the same as in the first null model. This is crucial, because, as we show in the Supplementary Material, computation of projection matrices used for calculating standard errors rely on this design matrix. The two different design matrices encapsulate different assumptions made on the distributions of the residuals. In the Supplementary Material, we show that in the absence of rank-normalization, a two-stage procedure in which (raw) residuals are used without covariate adjustment can lead to a loss of power. This is true when using rank-normalized residuals as well: to see this, assume that the residuals from the first stage regression are in fact normally distributed. Then rank-normalization has no effect, and our mathematical derivation demonstrates this.

### GWASs in the Hispanic Community Health Study/Study of Latinos

To study the effect of using residuals with and without adjusting for covariates, and with and without applying rank-normalization on the residuals, we performed multiple GWASs for each of 19 traits using up to 12,595 individuals from the Hispanic Community Health Study/Study of Latinos (HCHS/SOL). All participants provided informed consent and the study was approved by IRBs in each of the participating institutions. All models were adjusted to age, sex, field center, background group, log-transformed sampling weights, and the five first principle components representing distant genetic ancestry, and some traits were adjusted to additional covariates, such as age^2^, and BMI. We used linear mixed models with correlation matrices corresponding to community, household, and kinship (estimated from the genetic data). Information about the HCHS/SOL, genotyping, imputation, and genetic analysis in the HCHS/SOL, are provided in the Supplementary Material. Genotype and imputed data of the HCHS/SOL can be requested via dbGaP study accession phs000880. Phenotype data can be requested via dbGaP study accession phs000810.

### TOPMed hemoglobin WGS association study

We used data from TOPMed Freeze 4, which included 7,486 individuals from the Old Order Amish Study (N=1,102), Jackson Heart Study (JHS, N=3,251 African Americans), and Framingham Heart Study (FHS, N=3,133 European Americans from the offspring and generation 3 studies). All participants provided informed consent and the study was approved by IRBs in each of the participating institutions. Additional information about these studies is provided in the Supplementary Material. For TOPMed WGS data acquisition and QC report see ncbi.nlm.nih.gov/projects/gap/cgi-bin/GetPdf.cgi?id=phd006969,1.

We compared the performance of the SKAT Score statistic for testing the association of hemoglobin (HGB) values with sets of rare variants, under different approaches for rank-normalization and use of residuals, as described below. All analyses used linear mixed models, accounting for genetic relatedness using a genetic relationship matrix (GRM) computed using the GCTA method, (ref.15) where GRM was computed using all variants with MAF ≥ 0.001. Covariates were age, sex, and study. Additionally, to account for heteroscedasticity, we used a study-specific variance model, where we estimated a separate residual variance for each study (Amish, JHS, FHS Offspring, and FHS generation 3). The SKAT test was applied on sets of genotypes formed by taking all genetic variants with alternate allele frequency in the range (0,0.01), and dividing them into non-overlapping sets, defined by running windows across the genome, of length 5, 10, and 50 kilo bases (kb). For comparison, we also report results from analysis of a single permutation phenotype. Specifically, we randomly permuted HGB across participants once, and performed the same association testing as for the unpermuted trait.

### GWAS and WGS association studies – model comparisons

For genetic analyses in both the HCHS/SOL and TOPMed, we consider the approaches described in Table 1 and Figure S1 in the Supplementary Material. Table 1 describes the steps taken in each of the two stages (or in a single stage, for one of the approaches). For a given dataset, the covariates (when used) were always the same, as well as the GRM and variance component structure.

**Table 1:** analysis approaches compared in simulations and applied data analysis. For each of the compared analysis approaches (left column) the table provides the regression models from the two (or single) stages. Stage 1 is the same for all two-stage approaches. The association tests are general, and could be a single variant Wald, or a variant-set SKAT test, depending on the application.

| Approach | Stage 1 | Stage 2 |
| --- | --- | --- |
| Resid-Adj | Regress $Y \sim X$, giving<br>$\epsilon = Y - X\hat{\beta}$ | Test G association based on the regression $\epsilon \sim X + G$ |
| Resid-Unadj | | Test G association based on the regression $\epsilon \sim G$ |
| RN-Resid-Adj | | Test G association based on the regression $RN(\epsilon) \sim X + G$ |
| RN-Resid-Unadj | | Test G association based on the regression $RN(\epsilon) \sim G$ |
| RN-Trait | Test G association based on the regression $RN(Y) \sim X + G$ | -- |

All null models and subsequent association tests were computed in R using the GENESIS package (ref.16). For the HCHS/SOL, we focused on a single-variant Wald test, calculating the p-values of test statistics based on the *N*(0,1) distribution using model-based standard errors, and for TOPMed, we focused on a variant-set SKAT test (ref. 17). To quantify inflation, we used the inflation factor *λ_gc_* (ref.18) computed as the ratio between the quantile of the 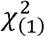 distribution corresponding to the median observed p-value ***p_med_***, computed as 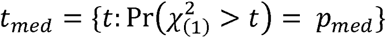 and the median value of the 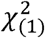 distribution, computed as 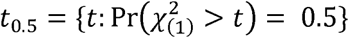, i.e. *λ_gc_* = *t_med_/t*_0.5_.

### Simulation studies

We performed a simulation study mimicking the settings in (ref.12), with modification to examine the effect of covariate confounding by a genetic principal component. We generated outcomes according to the model *y_i_* = *x_i_β_a_* + *g_i_β_g_* + *ϵ_i_*,*i* = 1,…*n*, where the residual *ϵ_i_* was sampled from three different distribution settings: ‘normal’ *ϵ_i_* ~ *N*(0,1); ‘outlier’ *ϵ_i_* ~ *aN(*0,1) + (1 − *a)N(*0,3), with *a* = 1 with probability 0.99, and 0 otherwise; or ‘non-normal’ 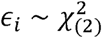 There was a single covariate *x_i_* ~ *N*(0,1), mimicking a single continuous measure of ancestry, with effect fixed at *β_a_* = 1. Genotype values were simulated under varying minor allele frequencies, and with and without association to the covariate, as follows. A genotype value for each person was sampled in Hardy-Weinberg Equilibrium, i.e. from a binomial distribution, with *g_i_~ Binom*(*p_i_*, 2), and probability computed as *p*_i_ = *exp*(*γ*_0_ + *x_i_γ_x_*)/(1 + *exp*(*γ*_0_ + *x_i_γ_x_*), with *γ*_0_ ϵ {−2, −3, −4, −5, −6, −7} controlling how rare the minor allele is. At the average value of ancestry, i.e. *x_i_* = 0, the lowest value *γ_0_* = –7 gives MAF=0.0009, and the highest value *γ_0_* = –2 gives MAF =0.12. Lastly, *γ_x_* determines the strength of confounding induced by the ancestry covariate, with *γ*_0_ ϵ {0,1,2}, where *γ_x_* = 0 corresponds to no confounding. While this model is consistent with confounding due to ancestry, other mechanisms in which the genotype is associated with the covariate (e.g. BMI, age in a cohort of old individuals) are also possible and have the same downstream effect on association analyses. We performed 10^7^ simulations from each combination of parameter settings and *β* = 0 (under the null), enabling a good estimation of type 1 error rate at the 10^-4^ significance level. Type I error was computed as the proportion of simulations in which the null hypothesis was rejected (p-value <10^-4^). For power, we ran 10^4^ simulations for each combination of the aforementioned parameters and with *β* ϵ {0.1,0.15,0.2). Power was generally calculated as the proportion of simulations in which the null hypothesis was rejected. In instances in which the Type I error was not control, we calculated the significance level yielding the desired Type I error rate, and used this threshold when determining whether the null hypothesis is rejected.

## Results

### Simulation studies

Comprehensive simulation results are provided in the Supplementary Material. We here provide a summary of the results, together with selected figures that demonstrate the main results for Type I error rate control and power.

In the ‘normal’ simulation settings (normally distributed errors), all tests always controlled Type I errors appropriately (Supplementary Material, Figures S5a and S6a). Figure 1 provides power estimates for these settings, for varying degrees of confounding and for common (MAF ranged between 0.11 to 0.22) and rare (MAF ranged between 0.002 to 0.02) variants. The power of the unadjusted methods (RN-Resid-Unadj, Resid-Unadj) decreases, compared to the adjusted methods (RN-Resid-Adj, Resid-Adj, RN-Trait-Adj) as the confounding effect increases. All adjusted methods are roughly similar for common variants, however, RN-Trait-Adj loses power compared to Resid-Adj and RN-Resid-Adj when variants become rare. Complete power figures for this setting are provided in the Supplementary Material, Figures S5b and S6b.

**Figure 1:**
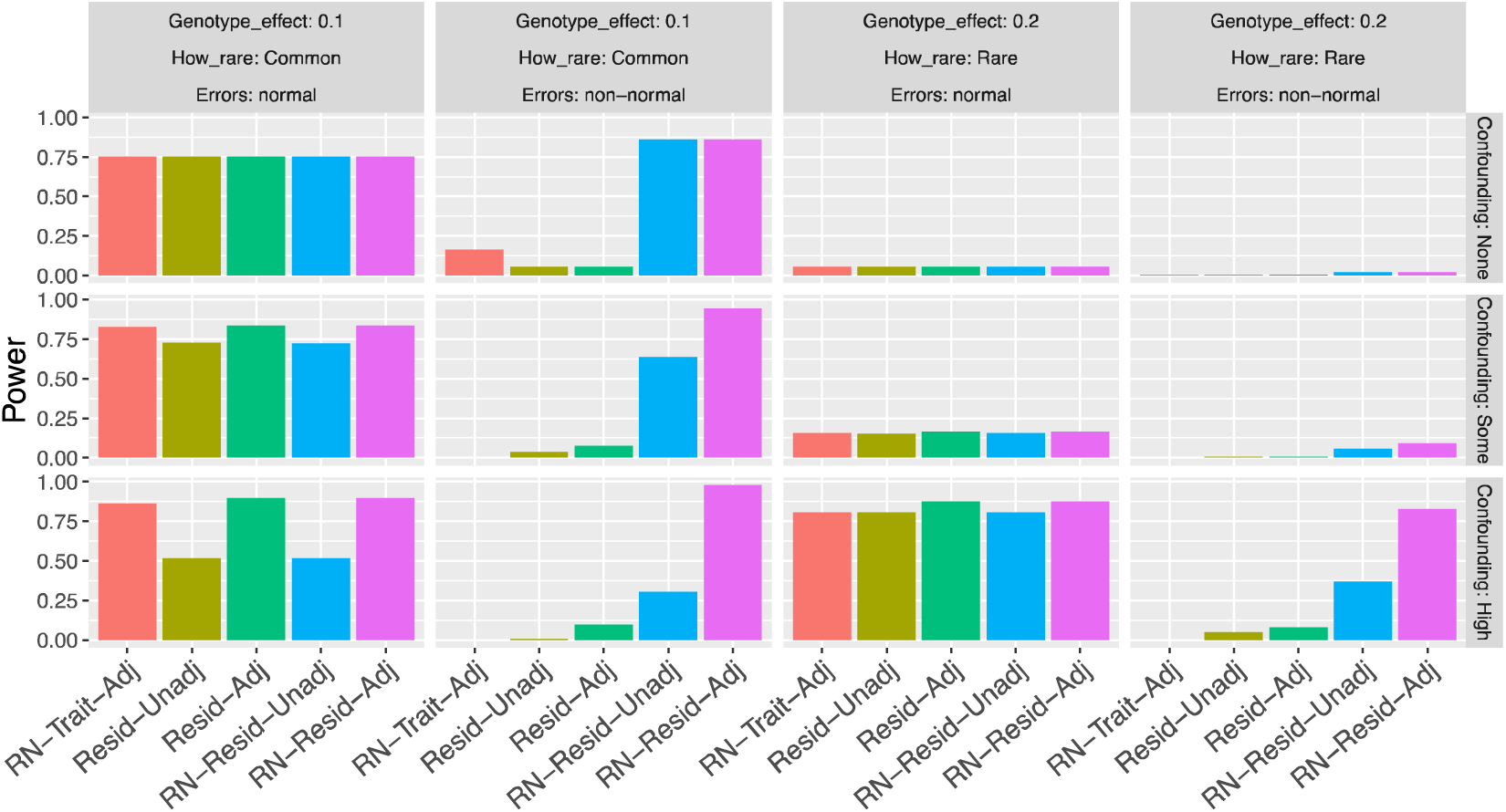
Estimated power in selected simulation studies. Each of the panels provides the power calculated over 10^4^ replicates of simulations, for each of the compared analysis approaches. Here we consider the ‘normal’ and ‘non-normal’ distribution settings, focusing on rare and common variants (Common: *γ_0_* = –2; Rare: *γ_0_* = –5) and by varying degrees of covariate confounding (None: *γ_x_* = 0; Some: *γ_x_* = 1; High: *γ_x_* = 2). In the displayed results, sample size was n=10,000, p-value threshold for determining significance was set at 10^-4^.

Selected results from the ‘non-normal’ simulation settings (chi-squared distributed errors) are presented in Figure 1 (power) and Figure 2 (type 1 error rates). Type I errors are well controlled when variants are common and there is no confounding. However, alarmingly, Type I error rates are very high for common variants when there is confounding effect, for RN-Resid-Unadj and RN-Trait-Adj. As variants become rare, all methods do not control Type I error under some conditions. Moreover, for rare variants, methods that do not rank-normalize (Resid-Adj and Resid-Unadj), did not control the Type I error rate even under no confounding. While RN-Resid-Adj did not control Type I error rate in the rare variant and strong confounding settings, overall it was the “least bad” approach of those investigated, with lowest observed levels of inflation overall. Figure 1 provides (calibrated) power for the ‘non-normal’ settings. Here, in the common variant and no confounding scenario, where all methods controlled the Type I error rate, only RN-Resid-Adj and RN-Resid-Unadj had high power. The pattern was similar across settings (see Figures S7 and S8 in the Supplementary Material for complete results).

**Figure 2:**
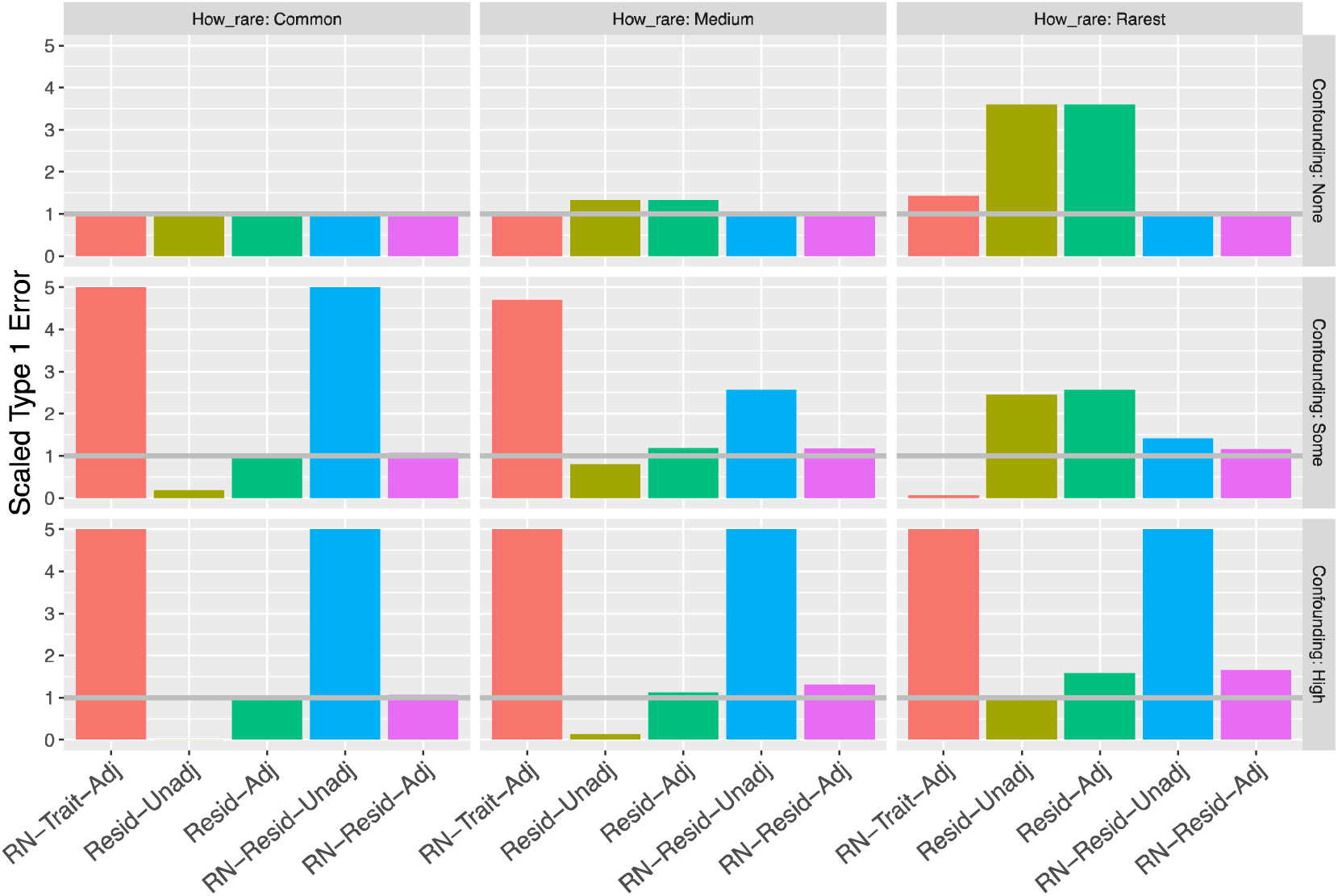
Type 1 error rates in the ‘non-normal’ simulation settings. Each of the panels provides the (scaled) estimated type 1 error rate over 10^7^ replicates of simulations, respectively, for levels of variant frequency (Common: *γ_0_* = –2; Medium: *γ_0_* = –5; Rarest: *γ_0_* = –6), and degree of confounding (None: *γ_x_* = 0; Some: *γ_x_* = 1; High: *γ_x_* = 2). Type 1 errors are scaled by the expected type 1 error rate (ideal value is 1, higher values indicate high rate of false positives, or inflation, lower values indicate deflation, or conservatism). In the displayed results, sample size was n=10,000, p-value threshold for determining significance was set at 10^-4^.

Finally, the results from the ‘outlier’ simulation settings (errors from a mixture of two normal distributions with different variances) had intermediate results compared to the two more extreme cases of ‘normal’ and ‘non-normal’ simulations. Results from these settings are provided only in the Supplementary Material. The Type I error rate was inflated under some conditions, with, for a given analysis approach, inflation generally higher as a variant becomes rarer and for smaller sample sizes (Supplementary Material Figures S9a and S10a), and with inflation being better controlled with rank-normalization, and power being higher for fully adjusted methods under confounding effect (Supplementary Materials Figures S9b and S10b).

### GWASs in the HCHS/SOL

We report results from HCHS/SOL GWASs of 19 traits in the Supplementary Material, and here highlight analyses of 4 traits: height, systolic blood pressure (SBP), ferritin, and number of teeth (N-teeth), in Figure 3. For all traits, Figures S11-S30 in the Supplementary Material demonstrate that Resid-Adj is always had smaller p-values than Resid-Unadj, in agreement with the derivation in the Supplementary Material. Similar patterns are observed when comparing RN-Resid-Adj to RN-Resid-Unadj when the trait residuals are relatively normally distributed. For example, the top left panel in Figure 3 compares the p-values from the GWASs of height in using RN-Resid-Adj and RN-Resid-Unadj, and the patterns is essentially the same as that seen in the analyses without rank-normalization, Resid-Adj and Resid-Unadj (see Figure S13 in the Supplementary Material for distribution of height residuals). When traits are further from normalilty (SBP, ferritin, N-teeth), this pattern changes, and we see some genetic variants with lower p-value in RN-Resid-Unadj compared to RN-Resid-Adj (Figure 3). The most extreme example is N-teeth, for which *λ_gc_* was 1.02 and 1.77 with and without second-stage adjustment for covariates, respectively, i.e. RN-Resid-Unadj was highly inflated. These results are in line with the simulations provided in Figure 2, where in the ‘non-normal’ setting, even a relatively small degree of confounding caused a large inflation of RN-Resid-Unadj.

**Figure 3:**
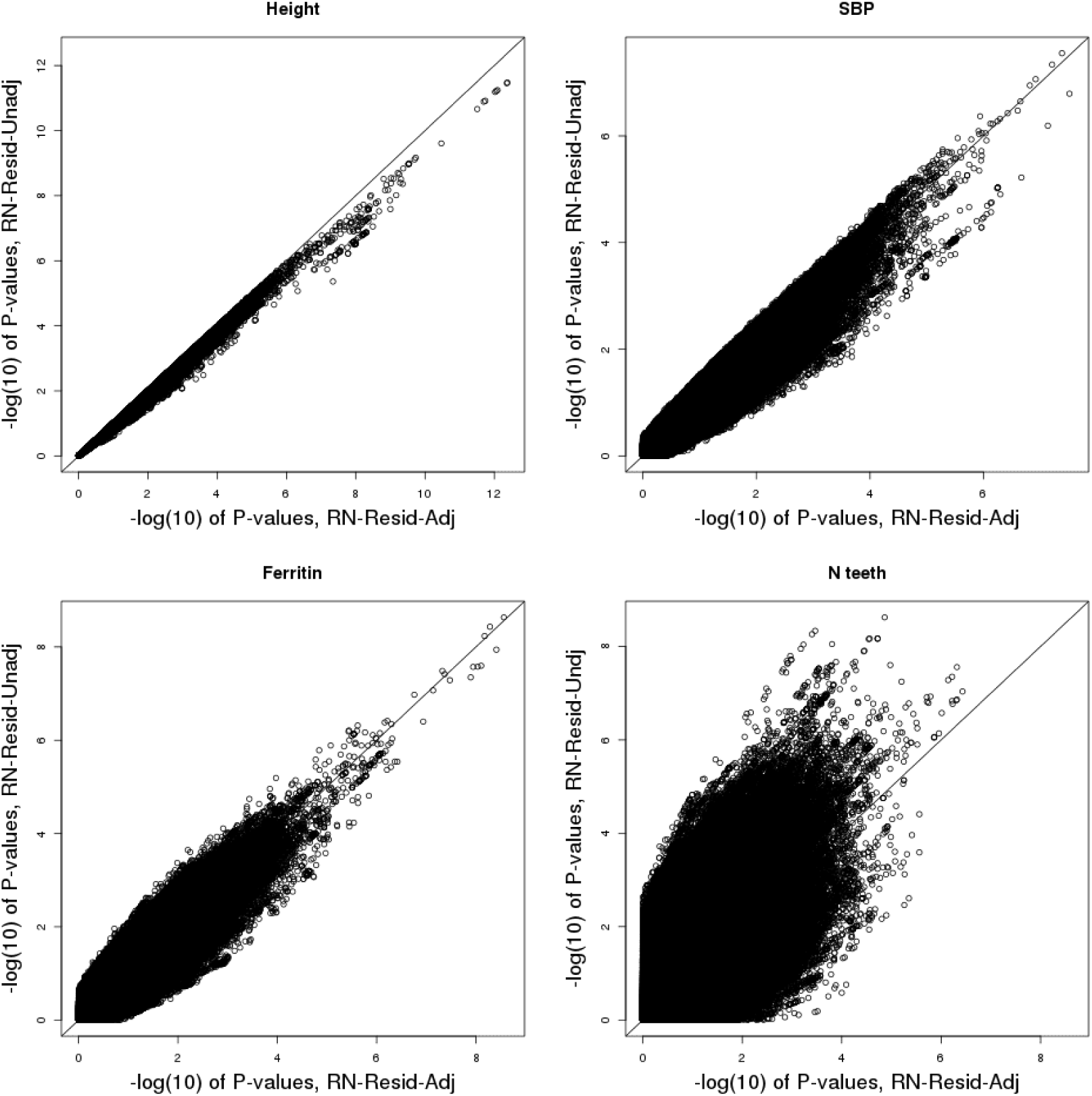
Computed p-values from the analyses of height, SBP, ferritin, and N-teeth in participants of the HCHS/SOL. Each of the panels corresponds to a different trait, and compares the (–log) p-values obtained from analyses that tests the association between rank-normalized transformed residuals and common genotypes (MAF≥ 0.05), with and without adjustment to the covariates that were used in the model that obtained the residuals.

### Hemoglobin WGS study in TOPMed

Figure 4 compares the inflation factors across scenarios and sliding window sizes, under true and randomly permuted HGB concentrations. Using Resid-Unadj and RN-Resid-Unadj produced deflated results, compared to Resid-Adj and RN-Resid-Adj, across all settings, as expected. In both real and permuted HGB, the difference in inflation factors between covariates adjusted and unadjusted settings is larger when the residuals were not rank-normalized; and the *λ_gc_* values are about the same in RN-Trait and RN-Resid-Adj settings. In terms of distribution, as seen in Figure S1 in the Supplementary Material, HGB residuals have a few negative outlying values, but otherwise their distribution is relatively normal. This is likely why we primarily see less significant p-values in the unadjusted analyses.

**Figure 4:**
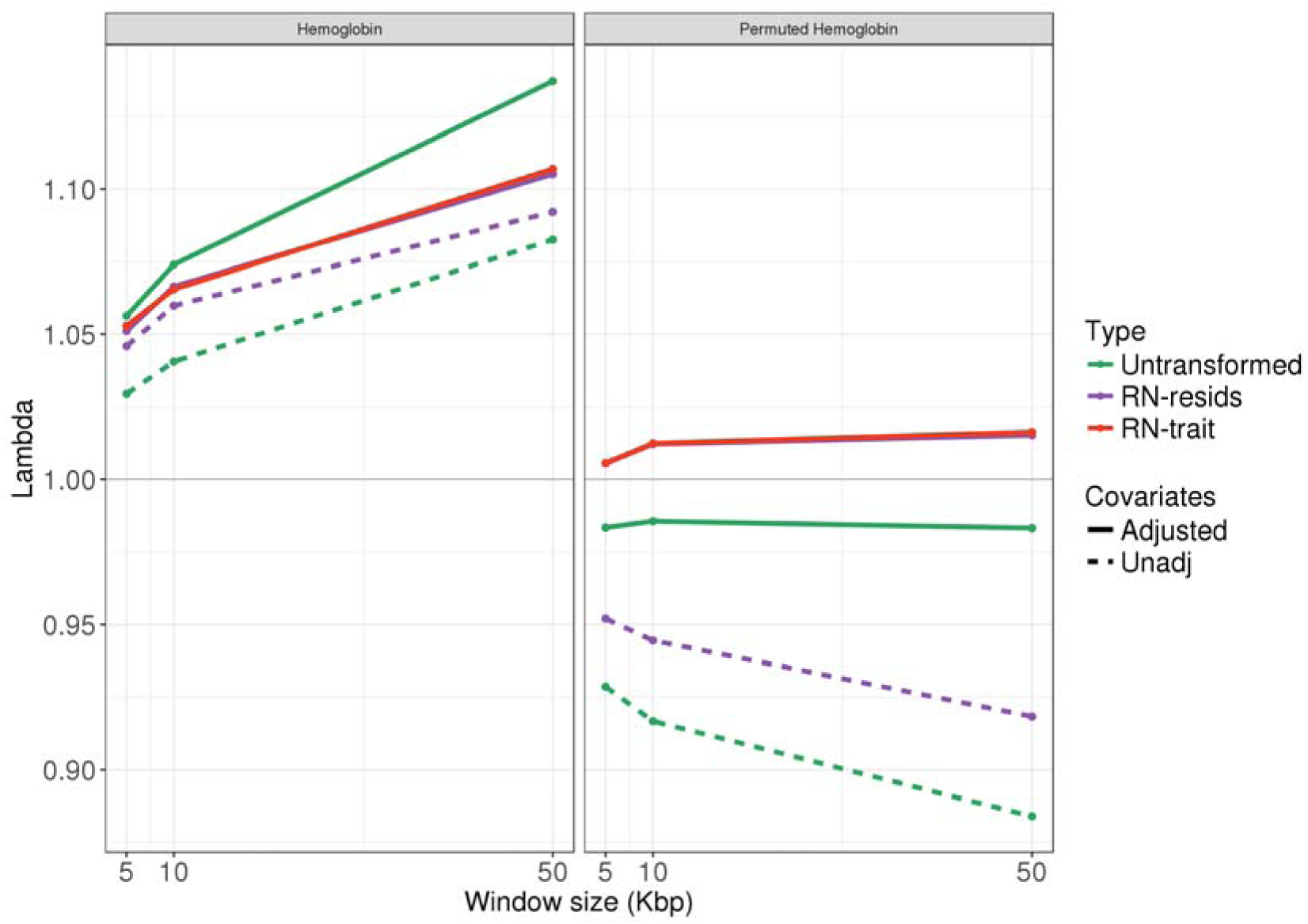
Observed inflation factors in SKAT analyses of the TOPMed Hemoglobin dataset. The figure provides the observed inflation factors lambda = *λ_gc_* in testing variants with alternate allele frequencies between 0 to 0.01 in non-overlapping window of sizes 5,10, and 50 Kbp. The left panel corresponds to the true hemoglobin trait, and the right panel correspond to the same analyses applied on permuted values of the observed hemoglobin.

In the Supplementary Material, Figures S2-S4, we provide p-values comparisons for the SKAT tests results obtained using the Resid-Adj, Resid-Unadj, RN-Resid-Adj and RN-resid-Undj. Using residuals without rank-normalization produced very low p-values for some of the tested variant sets. For these sets, there was no difference between Resid-Adj and Resid-Unadj, matching the pattern in Figure 2 (non-normal outcome) under the rarest variant and no confounding, where rank-normalization helped control inflation.

## Discussion

The validity of linear regression-based tests of the genetic association with a trait can be sensitive to the trait’s distribution. Some analysis approaches that have been used to counteract this problem include rank-normalization of both trait values and of residuals, which are then used in a partly-adjusted two-stage procedure. Both approaches are known to suffer from drawbacks, albeit some investigators have argued that they are appropriate in the context of rare variants analysis (ref.4). Here we have proposed a fully-adjusted two-stage approach, which uses the rank-normalized residuals as the outcome in the genetic association testing stage – a stage in which we again adjust for the same covariates used in the first stage. This approach ameliorates the problems of previous methods, for analysis of both common and rare variants. However, as we show in simulations, under non-normality of the trait and confounding by covariates, all the tests of low-count variants we considered may be biased. We separated the roles of adjustment for covariates and rank-normalization and showed theoretically (in the Supplementary Material) that, without rank-normalization, an unadjusted two-stage procedure may result in loss of power, when using either Wald or Score tests. We demonstrated the shortcomings of the partly-adjusted two-stage procedure in both a GWAS, interrogating common variants, and in a WGS study, testing rare variants.

For common variants, previous criticism of the partly-adjusted two-stage approach showed that it results in biased effect size estimates when covariates confound the genotype-trait associations, (ref.10) and that it loses power (ref.10, 11) compared to a one-stage approach testing the trait directly. The fully-adjusted two-stage approach alleviates both of these concerns: including covariates in the second stage alleviates the confounding problem, because these confounders are accounted for. This is demonstrated in the data analysis example from the GWASs in the HCHS/SOL. We first saw, for all GWASs, that a two-stage procedure of the form Resid-Unadj loses power compared to Resid-Adj, which recovers the same results from the untransformed trait-based analysis. When we applied rank-normalization to the residuals (RN-Resid), we saw that under non-normality, the confounding effects reported by (ref.10) in action, so that some of the unadjusted two-stage procedure RN-Resid-Unadj GWASs had many highly significant findings, which are likely false positives, compared to the fully adjusted procedure RN-Resid-Adj, with N-teeth GWASs being the most extreme example.

For rare variants, previous work by Tang and Lin (ref.4) showed in the context of meta-analysis that rank-normalizing the trait in one-stage analysis is useful. However, when pooling multiple heterogeneous studies together in a joint analysis, as in TOPMed, there are strong confounders (e.g., study), so the problems raised by Beasley et al. (ref.3), pointing at biases when residuals are non-normal and there is covariate confounding, are expected, and are demonstrated in our simulation study, in which RN-Trait-Adj analysis did not control Type I error rates. We note that our simulations show that when the trait is highly non-normal and covariate confounding of the genotype-trait association exists, all analysis methods may be inflated, including RN-Trait-Adj. However, RN-Resid-Adj had generally a lower degree of inflation compared to other methods.

Recently, Auer et al. (ref.12) argued that an unadjusted two-stage procedure is appropriate for rare variants, because the confounding problem pointed out in (ref.10) is negligible. In our simulations, we see that RN-Resid-Unadj controlled type 1 error in the normal and outlier outcome settings, where it lost some power compared to RN-Resid-Adj in the presence of confounding. In the analysis of TOPMed hemoglobin data set, we tested sets of rare variants using SKAT. Rank-normalizing either trait or residuals reduced overall inflation. Not adjusting for covariates in a non-adjusted two-stage procedure clearly caused a strong deflation, as measured by inflation factors, i.e. in the median of the distribution. For very low p-values, they remain qualitatively quite similar in the partly- and fully-adjusted two-stage procedures (Figures S3-S6 in the Supplementary Material). Still, the overall distribution of results is important and serves as a primary tool in evaluating model fit in genetic association studies. Therefore, one should be cautious about using a partly-adjusted approach.

In meta-analysis of GWASs, investigators often apply genomic control (ref.18) on each individual study contributing to the meta-analysis. This differs from our approach, in that genomic control assumes that there is global dispersion in the study, which affects all tested variants in a similar way, and can be globally corrected by a single constant. Instead our method is primarily concerned with the effect of non-normality on association analysis. Non-normality affects different genetic variants in different ways. Similarly, confounding effects, as our GWAS results demonstrate, differ between genetic variants. Therefore, a single global correction cannot account for, or fix, the challenges that we highlighted.

In summary, in this investigation we provide a thorough assessment of the controversial uses of the rank-normalizing transformation which is often used in practice despite several published manuscripts criticizing their use. We demonstrate a proper and beneficial use of such transformations when coupled with a fully adjusted two-stage procedure. In addition to the main investigation, in the Supplementary Material (Figures S12-S30) we provide comparisons of the approaches investigated in this manuscript for GWASs of 19 anthropometric, blood pressure, blood markers, and electrocardiogram traits in the HCHS/SOL, alongside the distribution of their residuals from the ‘null model’. These comparisons suggest that future large consortia meta-analyses may reduce Type I errors and gain power from using the fully-adjusted two-stage approach, compared to the partly-adjusted approach often used.

Supplementary information is available at European Journal of Human Genetics’ website.

## Acknowledgements

TS was supported by NHLBI R01HL120393-03S1 and 1R35HL135818, and NHGRI R01HG005827. LMR was supported by T32 HL129982.

## References

1. Wu X, Cooper RS, Borecki I, Hanis C, Bray M, Lewis CE, et al. A combined analysis of genome-wide linkage scans for body mass index from the National Heart, Lung, and Blood Institute Family Blood Pressure Program. Am J Hum Genet. 2002;70(5): 1247–56.

2. Ashton GC, Borecki IB. Further evidence for a gene influencing spatial ability. Behav Genet. 1987;17(3):243–56.

3. Beasley TM, Erickson S, Allison DB. Rank-based inverse normal transformations are increasingly used, but are they merited? Behav Genet. 2009;39(5):580–95.

4. Tang ZZ, Lin DY. Meta-analysis for Discovering Rare-Variant Associations: Statistical Methods and Software Programs. Am J Hum Genet. 2015;97(1):35–53.

5. Hoffmann TJ, Ehret GB, Nandakumar P, Ranatunga D, Schaefer C, Kwok PY, et al. Genome-wide association analyses using electronic health records identify new loci influencing blood pressure variation. Nat Genet. 2017;49(1):54–64.

6. Lange LA, Hu Y, Zhang H, Xue C, Schmidt EM, Tang ZZ, et al. Whole-exome sequencing identifies rare and low-frequency coding variants associated with LDL cholesterol. Am J Hum Genet. 2014;94(2):233–45.

7. Shungin D, Winkler TW, Croteau-Chonka DC, Ferreira T, Locke AE, Magi R, et al. New genetic loci link adipose and insulin biology to body fat distribution. Nature. 2015;518(7538):187–96.

8. Tajuddin SM, Schick UM, Eicher JD, Chami N, Giri A, Brody JA, et al. Large-Scale Exome-wide Association Analysis Identifies Loci for White Blood Cell Traits and Pleiotropy with Immune-Mediated Diseases. Am J Hum Genet. 2016;99(1):22–39.

9. Wen W, Cho YS, Zheng W, Dorajoo R, Kato N, Qi L, et al. Meta-analysis identifies common variants associated with body mass index in east Asians. Nat Genet. 2012;44(3):307–11.

10. Demissie S, Cupples LA. Bias due to two-stage residual-outcome regression analysis in genetic association studies. Genet Epidemiol. 2011;35(7):592–6.

11. Che R, Motsinger-Reif AA, Brown CC. Loss of power in two-stage residual-outcome regression analysis in genetic association studies. Genet Epidemiol. 2012;36(8):890–4.

12. Auer PL, Reiner AP, Leal SM. The effect of phenotypic outliers and non-normality on rare-variant association testing. Eur J Hum Genet. 2016;24(8):1188–94.

13. McCullagh P, Nelder JA. Generalized linear models. 2nd ed. Boca Raton: Chapman & Hall/CRC; 1998. xix, 511 p. p.

14. Wu MC, Lee S, Cai T, Li Y, Boehnke M, Lin X. Rare-variant association testing for sequencing data with the sequence kernel association test. Am J Hum Genet. 2011;89(1):82–93.

15. Yang J, Lee SH, Goddard ME, Visscher PM. Genome-wide complex trait analysis (GCTA): methods, data analyses, and interpretations. Methods Mol Biol. 2013;1019:215–36.

16. Conomos MP, Thornton T, Gogarten SM, Brown L. GENESIS: GENetic EStimation and Inference in Structured samples (GENESIS): Statistical methods for analyzing genetic data from samples with population structure and/or relatedness. 2.8.0. ed. Bioconductor, https://bioconductor.org/packages/release/bioc/html/GENESIS.html2017.

17. Lee S, Miropolsky L, Wu M. SKAT: SNP-Set (Sequence) Kernel Association Test. R package version 1.3.2.1 ed 2017.

18. Devlin B, Roeder K. Genomic control for association studies. Biometrics. 1999;55(4):997–1004.

